# Patterns and predictors of human-sloth bear conflict in Nepal: Implications for coexistence in unprotected forest landscapes

**DOI:** 10.1101/2022.08.24.505134

**Authors:** Manoj Pokharel, Chandramani Aryal, Bidhan Adhikary, Jeevan Rai

**Author notes:** Corresponding author: Manoj Pokharel Nature Conservation and Study Centre, P.O. Box: 2175 Kathmandu, Nepal.

## Abstract

Human-sloth bear conflict, generally in the form of human attacks, is common throughout most areas where sloth bears co-occur with humans. The situation is more prevalent in multi-use forest landscapes outside protected areas. Although sloth bears are a conflict-prone species in Nepal, there is a clear lack of systematic studies that can inform human safety and conflict mitigation. We used data from questionnaire-based interviews with conflict-affected people and witnesses to provide detailed information about human-sloth bear conflict (1990– 2021) in the Trijuga forest, an important sloth bear stronghold outside protected areas in Nepal. The data were analyzed using descriptive statistics, chi-square tests, and regression analysis. For the time period, 66 conflict incidents involving 69 human individuals were recorded, with an annual average of 2.06 (SD = 1.48) incidents and 1.75 (SD = 1.34) attacks. Conflicts primarily impacted working-age group (25–55 years old) men, whose primary occupation was farming and who frequented the forest regularly. They typically occurred between 0900 and 1500, inside forests, and in habitats associated with poor land cover visibility. Poor visibility was also a significant positive determinant of bear attacks on humans. Fifty-six conflict incidents resulted in attacks that injured 59 people, with a fatality rate of 8.47%. Victims of bear attacks frequently had serious injuries, especially to the head and neck areas of the body. Serious injuries were more likely to occur to lone individuals than to people who were in groups of two or more. We suggest identification of conflict-risk habitats through a participatory mapping approach and education programs for the local people for effective human-sloth bear conflict management in Nepal’s unprotected forests.

## Introduction

Large carnivores’ through the trickle-down control approach, play an important role in shaping the structure and function of ecosystems that they inhabit [1–3]. Owing to their position on food chain, they need larger area to fulfill their ecological requirements, increasing the chances for encounter and interaction with the humans [4]. As a result, large carnivore management is a major priority in today’s conservation scenario, particularly in human-dominated settings where carnivore populations are diminishing [5]. Large carnivore management in human-dominated environments, on the other hand, poses a slew of problems, the most pressing of which is arguably human-carnivore conflict [4,5]. Direct attacks on humans by large carnivores, among other types of conflict, can have serious and far-reaching implications for both carnivores and humans [6,7]. On the contrary, reliable information on the characteristics of such incidents might assist people in avoiding or minimizing such circumstances [8,9]. This, in turn, can help garner support from general public to protect and coexist with large carnivores, which is especially useful in areas where humans are present on a regular basis in carnivore habitats [9–11].

The sloth bear (*Melursus ursinus*), despite exhibiting no predatory or territorial behavior [12], has evolved into one of the most aggressive large carnivore species on the Indian subcontinent [13,14]. This development is most likely a protective adaptation for coexisting with large and seemingly dangerous mammals, such as the Asiatic elephant (*Elephas maximus*), Bengal tiger (*Panthera tigris*), one-horned rhinoceros (*Rhinoceros unicornis*), and common leopard (*Panthera pardus*) [12,14,15]. Because of the widespread human intrusion in sloth bear habitat [16], sloth bears’ violent defense behavior has become a severe problem for human safety, as humans are routinely attacked, injured, or killed by sloth bears [17–19]. As a consequence, many people are afraid of sloth bears and often refuse to undertake or support conservation efforts that are aimed at them [16,20].

Researchers are increasingly looking into the features of human attacks by sloth bears in order to lessen the severity of the problem. However, the majority of such studies are only available from India or Sri Lanka [18–22]. In both countries, more intense conflicts with sloth bears have been reported from areas outside protection where disturbances are generally higher [18,20,23]. Given that site-specific natural and anthropogenic factors can bring about changes in conflict patterns involving sloth bears [19,20], researchers argue the need to study human-sloth bear conflicts in other range areas of the species to aid in the development of more effective and site-specific conflict minimization measures [24].

Although sloth bears are a conflict-prone wildlife species in Nepal’s southern region [25–27], literatures are limited to provide comprehensive information that can be utilized to inform human safety and conflict mitigation. The majority of recent studies focus on conflicts with sympatric species, such as tigers and elephants [28–31], and these are generally concentrated around protected areas [32]. Sloth bears receive little benefit from conflict mitigation implications derived from these studies due to their unique ecology and limited areas of distribution under protection. While the availability of conflict addressing measures and resources might have facilitated some level of coexistence between humans and sloth bears around Nepal’s protected areas [33], the unprotected areas lack such resources and therefore are more vulnerable to conflicts [34]. This necessitates evidence-based species-focused conflict mitigation measures that can be utilized by the local stakeholders in the unprotected areas [34].

By focusing on an important area for sloth bears outside of protection, this study aims to bridge the information gap on human-sloth bear conflict in Nepal and provide implications for conflict management and coexistence. Our objectives were to (1) document the socio-demographic characteristics of people affected by sloth bear conflicts, (2) determine the spatiotemporal patterns of conflicts, (3) investigate the characteristics and circumstances associated with conflicts, (4) categorize the injuries resulting from attacks, and (5) identify potential factors predicting attacks.

## Materials and Methods

### Study area

We carried out this study around the Trijuga (also known as ‘Triyuga’) forest situated in the southeast of Nepal (Figure 1). It is one of the largest remaining patches of Churia or Siwalik forest outside Nepal’s protected areas covering an area of approximately 430 km^2^ [35]. Available evidences indicate it to be the largest sloth bear stronghold outside protection in the country [25,36]. The forest is stretched east-west across the nine municipalities of the Udayapur and Saptari districts. It has a network of community forests across its periphery, while the central section is a part of national forest. *Shorea robusta* dominated dry deciduous forest predominates along the Trijuga forest’s northern boundary. Along the southern boundary mixed deciduous forest is prevalent with *Dalbergia latifolia*, *Acacia catechu*, *Terminalia tomentosa*, and *Semecarpus anacardium* as important species. The forest is bordered on most sides by agricultural lands and human settlements, resulting in a heavy reliance of the neighboring human communities for forest resources [36]. Apart from sloth bears, other major mammalian species found in this area include the common leopard, Asiatic elephant, barking deer (*Muntiacus muntjak*), wild boar (*Sus scrofa*), jungle cat (*Felis chaus*), golden jackal (*Canis aureus*), Bengal fox (*Vulpes bengalensis*), rhesus macaque (*Macaca mulatta*), Tarai gray langur (*Semnopithecus hector*), and Chinese pangoliln (*Manis pentadactyla*). The study area is described in more detail elsewhere [36].

**Figure 1:**
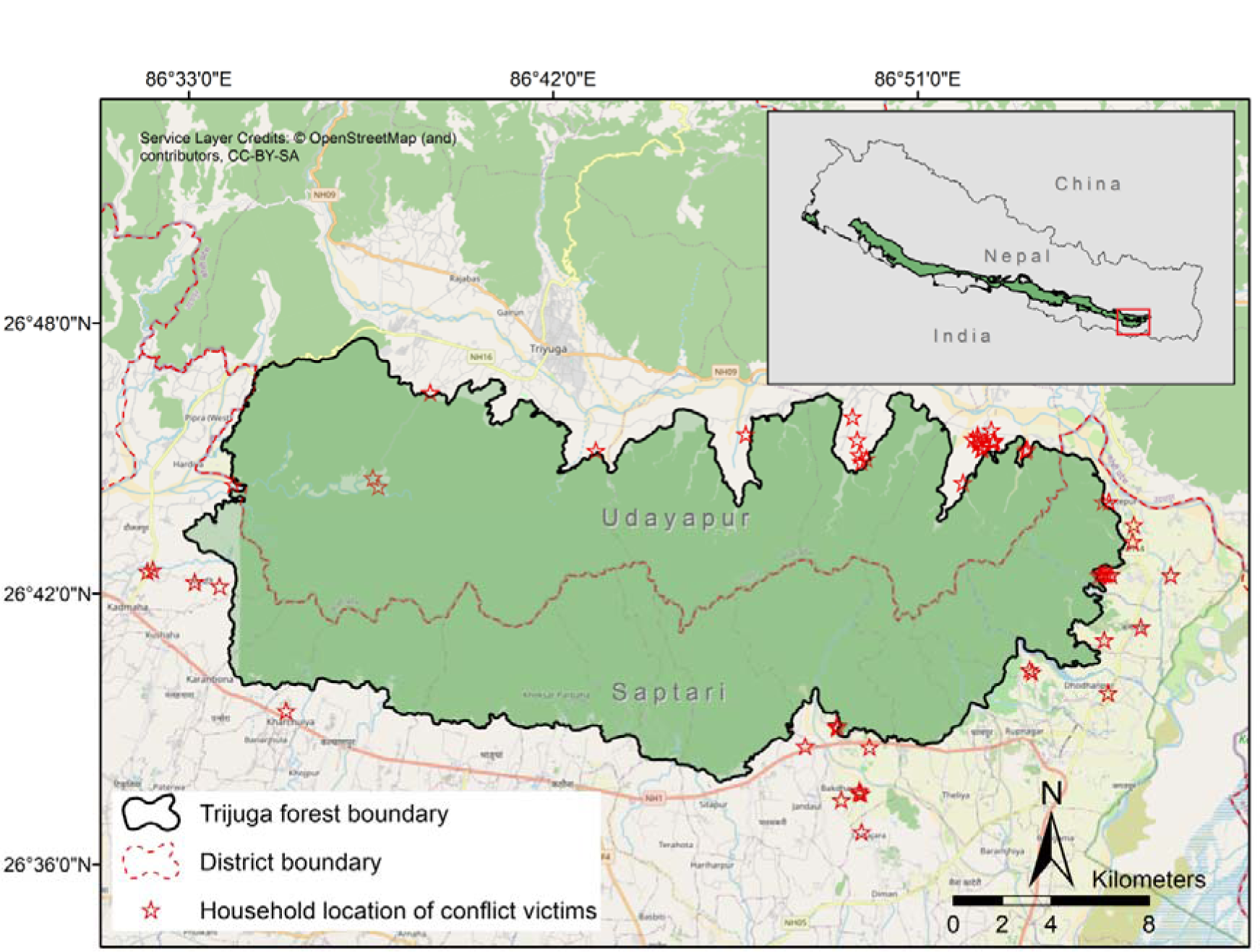
Map of Trijuga forest showing district boundary and household location of conflict victims. Inset map shows the location of the study area in reference to the predicted distribution range of sloth bears in Nepal, adapted from IUCN Red List 2020

### Data collection

In accordance with Hopkins et al. [37] vocabulary of terminology and concepts developed for human-bear study and management, we define human-sloth bear conflict as “any incident in which humans and bears come into close proximity, prompting the bear to react aggressively, resulting in the human taking evasive action and/or the bear making physical contact with the person”. Therefore, we also took into account the incidents where the people were not attacked but were at considerable risk of being attacked and injured by the bears [9,19].

Data on human-sloth bear conflict was collected between 15^th^ July and 23^rd^ September 2021 using a three-stage approach. In the first stage, we viewed records of relief applications or disbursements held by the forest offices, including division, sub-division and community forests. This provided us with a preliminary list of people around the Trijuga forest who had been affected by conflict with sloth bear. However, due to the Nepal government’s relatively recent initiation of relief mechanism for wildlife conflicts outside protected areas [34], and limited awareness among the local people about the relief mechanism, there was a high chance of conflicts going unreported to the authorities. Additionally, we were also interested in documenting unreported incidents where the encounters did not result in an attack but people were at significant risk of being attacked. In order to account for this, in the second stage we visited all the administrative divisions (2 Municipalities in Udayapur and 7 Municipalities in Saptari), that Trijuga forest is a part of, and spoke with the officials of local administrative bodies (Wards) and villagers to identify any remaining persons affected by sloth bear conflict. We set the upper limit for documenting conflicts at 1990 because we were able to identify only a small number of people (*n* = 3) who were involved in conflict before that year, and information provided for much older incidents was less likely to be reliable. In this way, we compiled a comprehensive catalogue of conflict victims from 1990 to 2021, and while the sampling strategy used was purposive, we believe it is representative of the study site for the time period.

In the final stage, we performed interviews with the conflict affected people, or in absence of such person, had closely witnessed the event (*n =* 10). A questionnaire containing a mix of closed-and open-format questions (Appendix 1) were used to collect data during the interviews [38,39]. The research methodology, including the questionnaire, was approved by the technical committee of the Department of Forests and Soil Conservation before issuing research permit. Approvals to carry out interviews were also obtained from the local administrative bodies. Before interviews respondents were made clear about the purpose of the study and verbal consent was obtained from each of them to start the interview [38]. Interviews were conducted in Nepali language, and in cases where the respondent could not speak or understand Nepali properly, a local field assistant helped in language translation.

The questionnaire was divided in two sections. The first section had questions asking respondents of: (a) temporal data (time, month and year of conflict); (b) spatial data (approximate spatial location, major land cover features and visibility conditions); (c) characteristics and circumstances associated with conflict (human and bear activity before encounter, human and bear behavioral response to encounter); (d) bear demographics (number, age, sex) and human companions (number, approximate distance) data; and (e) injuries from bear attacks (type of injuries and organs affected). The second section of questionnaire was focused in extracting socio-demographic information of conflict victims and had questions about the (a) ethnicity; (b) gender; (c) age; (d) occupation; (e) education level; (f) family size; (g) duration of residency; and (h) frequency and purpose of forest visit made by respondent, at the time of conflict. We also recorded household location of the respondents with the help of a Garmin Etrex 10 handheld GPS device.

### Data preparation and analysis

Microsoft Excel 2010 (Microsoft Redmond, USA) was used to enter and prepare the interview data for analysis. Five key categories were used to categorize respondents’ characteristics, including their age, ethnicity, and level of education (Table 1). The respondents’ occupation at the time of the conflict was divided into eight categories, and their frequency of visits to the forest was divided into four main classes (Table 1). Due to the rapid nature of the majority of human-sloth bear confrontations, it is acknowledged that human companions who are more than 50 meters away from a victim of a conflict have a relatively negligible effect on the outcome of the encounter [20]. Thus, the number of companions only included those who were within 50 meters of the victim. Likewise, it is known that land cover visibility plays a significant role in determining the occurrence of human-bear confrontations, including for sloth bears [9,24]. We followed Smith and Herrero [9] and Sharp et al. [24] qualitative classification of land cover visibility as: (1) poor (densely vegetated place or rough terrain with short sight distances, as well as periods of low light levels, such as dawn, dusk, or night); (2) moderate (areas with small patches of vegetation and limited tree cover allowing increased visibility than those rated poor); and (3) good (poorly vegetated open areas, such as wide sandy riverbanks and grasslands with short grasses).

**Table 1:**
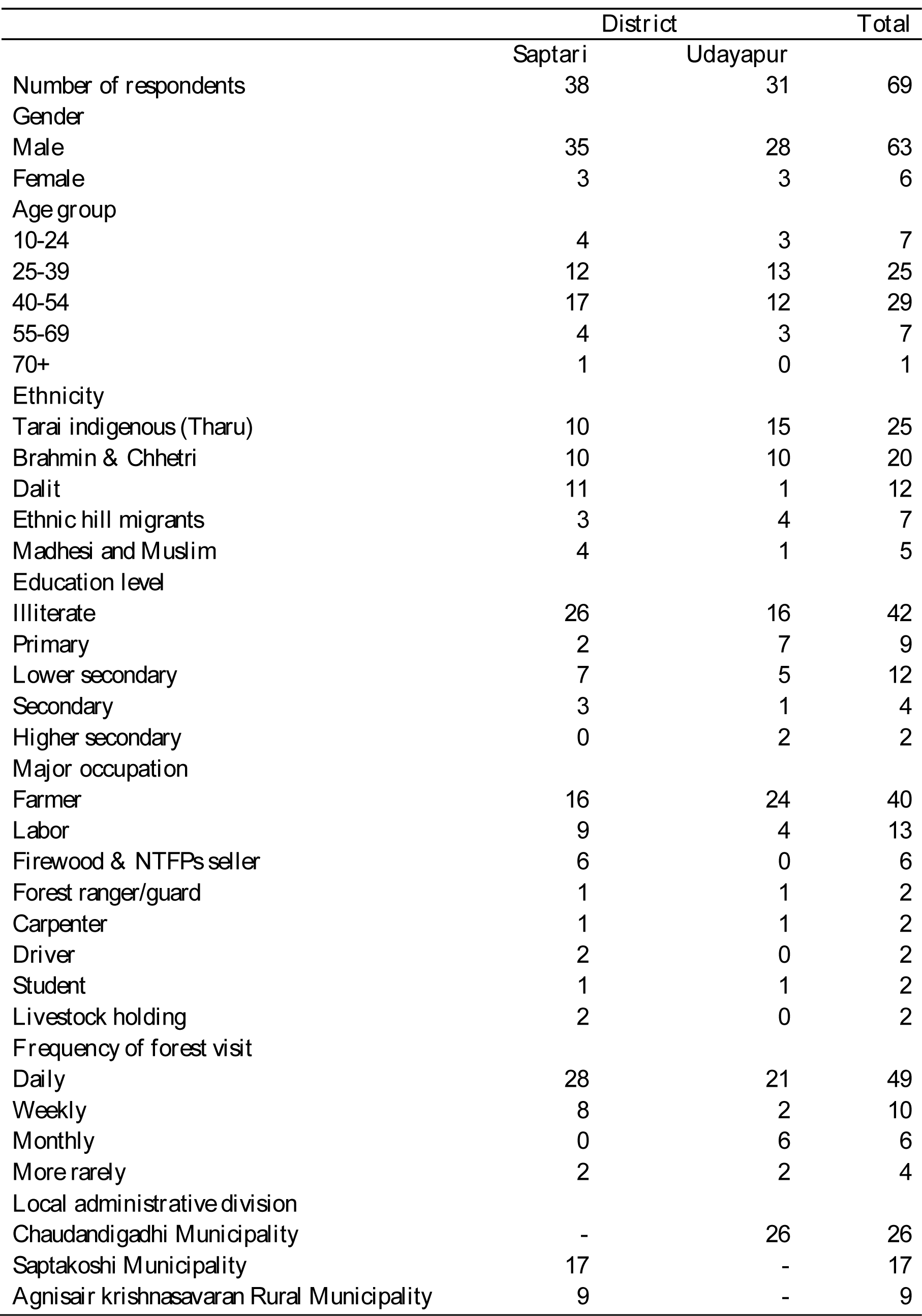
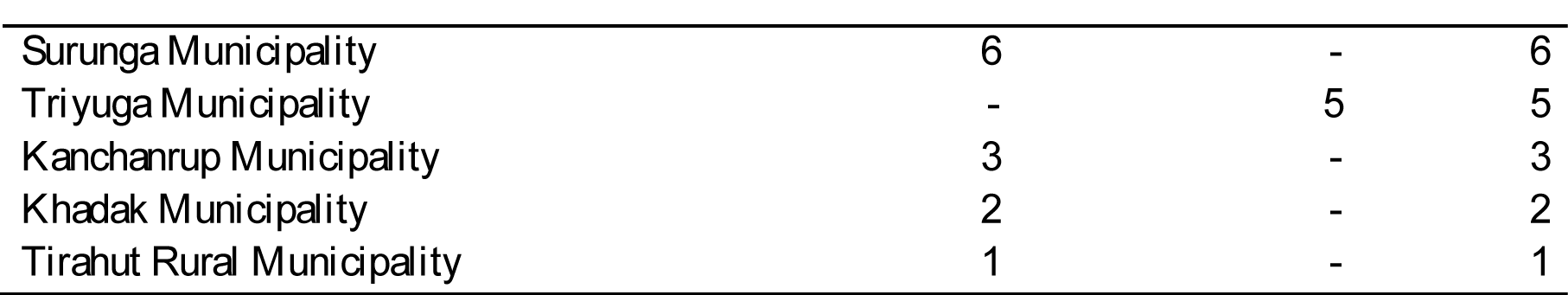
Socio-demographic profile of people affected by human-sloth bear conflict in the Trijuga forest from 1990 to 2021

|  | **District** |  | **Total** |
| --- | --- | --- | --- |
|  | **Saptari** | **Udayapur** |  |
| ***Number of respondents*** | 38 | 31 | 69 |
| ***Gender*** |  |  |  |
| Male | 35 | 28 | 63 |
| Female | 3 | 3 | 6 |
| ***Age group*** |  |  |  |
| 10-24 | 4 | 3 | 7 |
| 25-39 | 12 | 13 | 25 |
| 40-54 | 17 | 12 | 29 |
| 55-69 | 4 | 3 | 7 |
| 70+ | 1 | 0 | 1 |
| ***Ethnicity*** |  |  |  |
| Tarai indigenous (Tharu) | 10 | 15 | 25 |
| Brahmin & Chhetri | 10 | 10 | 20 |
| Dalit | 11 | 1 | 12 |
| Ethnic hill migrants | 3 | 4 | 7 |
| Madhesi and Muslim | 4 | 1 | 5 |
| ***Education level*** |  |  |  |
| Illiterate | 26 | 16 | 42 |
| Primary | 2 | 7 | 9 |
| Lower secondary | 7 | 5 | 12 |
| Secondary | 3 | 1 | 4 |
| Higher secondary | 0 | 2 | 2 |
| ***Major occupation*** |  |  |  |
| Farmer | 16 | 24 | 40 |
| Labor | 9 | 4 | 13 |
| Firewood & NTFPs seller | 6 | 0 | 6 |
| Forest ranger/guard | 1 | 1 | 2 |
| Carpenter | 1 | 1 | 2 |
| Driver | 2 | 0 | 2 |
| Student | 1 | 1 | 2 |
| Livestock holding | 2 | 0 | 2 |
| ***Frequency of forest visit*** |  |  |  |
| Daily | 28 | 21 | 49 |
| Weekly | 8 | 2 | 10 |
| Monthly | 0 | 6 | 6 |
| More rarely | 2 | 2 | 4 |
| ***Local administrative division*** |  |  |  |
| Chaudandigadhi Municipality | - | 26 | 26 |
| Saptakoshi Municipality | 17 | - | 17 |
| Agnisair krishnasavaran Rural Municipality | 9 | - | 9 |
| Surunga Municipality | 6 | - | 6 |
| Triyuga Municipality | - | 5 | 5 |
| Kanchanrup Municipality | 3 | - | 3 |
| Khadak Municipality | 2 | - | 2 |
| Tirahut Rural Municipality | 1 | - | 1 |

Since the date of conflict provided by the respondents was according to the Bikram Sambat calendar which is different from the Gregorian calendar, we used an online date converter tool (https://www.hamropatro.com/date-converter) to covert the dates to Gregorian calendar system. Further, the majority of respondents were able to recall the year and month but not the day of the conflict. To account for the fact that the Nepali months coincide with two English months (For example, the first month *Baishak* of the Bikram Sambat calendar runs from April 14 to May 14), we asked respondents whether the conflict occurred near the start, middle, or end of the Nepali month and selected the appropriate English month for the analysis. Based on the month of conflict we divided seasons as: (1) winter (December – February); (2) summer (March – May); (3) monsoon (June – September); and (4) autumn/post-monsoon (October – November).

To identify the approximate spatial location of the conflicts, we first divided the Trijuga forest into 4-km^2^ grid cells. We asked respondents for the name of the community forest (if applicable) where the conflict occurred, as well as the direction and distance to the conflict location from their home or from easily mapped features such as streams. In order to help respondents pinpoint the location of the conflict more precisely, we also showed printed satellite photographs of the study area that were obtained from Google Earth Imagery. Although exact coordinates could not be determined, the information provided for 58 conflicts was adequate to locate them within the 4-km^2^ grid cells. We mapped the number of conflicts according to grid cells and prepared a conflict hotspot map using ArcGIS 10.4.

In order to determine the pattern of injuries to human body from a bear attack, we divided the surface area of human body according to the Wallace’s “rule of nines” method [40] into six categories: (1) Head-neck region (9% of body surface); (2) back (18%); (3) front (chest & abdomen; 18%); (4) arms (9% each); (5) legs (18% each); and (6) perineum (1%). Further, we used criteria developed by Ratnayeke et al. [20], to determine the severity of injuries sustained during sloth bear attacks. These were grouped into four categories: (1) minor injuries (injuries that did not result in deep punctures and only had minor scratches or superficial lacerations); (2) moderate injuries (injuries that resulted in deep punctures that took longer to heal and left few permanent scars); (3) moderately serious injuries (fractures of the hands or feet, punctures to the head, several lacerations or punctures that resulted in significant bleeding); and (4) serious injuries (injuries that resulted in fractured bones in the head or neck, loss of scalp, the removal of facial components (such as the eyes, nose, and ears), or the permanent impairment of one’s vision, hearing, or speech.

We computed descriptive statistics (mean, standard deviation, and frequencies) to summarize the data. The disparity between observed and expected frequencies across various categories of a variable was examined using a Chi-square goodness-of-fit test. We specifically investigated whether conflict rates differed by ethnic groups, age groups, frequency of forest visits, year group, months, seasons, and time of the day. We also looked at whether the frequency of injuries varied according to the four severity classes and whether some body parts were more vulnerable to injuries than others. A chi-square test of independence was performed to determine if there is an association between human group size and the severity of the injuries sustained due to bear attacks. The strength of association was calculated using Cramer’s V. Significance level for all the chi-square analysis was determined at α ≤ 0.05.

Furthermore, we used non-parametric linear regression using the ‘Rfit’ package [41] in R to assess the trend of human-sloth bear conflict by taking year as an explanatory variable and the number of conflicts as a response variable. Additionally, we selected four variables (Table 2) that could potentially predict the outcome of a human-bear encounter (attack or non-attack, coded as 1 or 0, respectively). In order to assess the relative importance of variables we built regression models using the firth’s bias-reduced logistic regression through the use of R package ‘logistf’. Firth’s logistic regression effectively takes into consideration the problem of statistical separation and small sample size of the data [42]. Small sample size-corrected Akaike Information Criterion (AICc) was used for comparison and selection of the candidate models [43], using the R package ‘MuMIn’. Model selection uncertainty was accounted for by model averaging the competing models (ΔAICc ≤ 2), and estimates of regression coefficients (β estimates) were generated to determine the direction and magnitude of influence of the variables on the probability of bear attack.

**Table 2:** Variables devised to test their influence on the probability of a sloth bear attack

| Variable | Type | Description/categories |
| --- | --- | --- |
| Human group size | Numeric | Number of people within 50 m of the conflict victim |
| Bear group size | Numeric | Number of bears engaged in conflict |
| Land cover visibility | Categorical | Poor (yes) or not poor (no) |
| Making noise | Categorical | Victim or companion engaged in making noise before encounter (yes) or not (no) |

## Results

We documented 66 incidents of human-sloth bear conflict during a 32-year period (1990– 2021). Of the total, 56 events (84.84%) resulted in human attacks, while the remaining ten did not result in physical contact. The conflicts involved 69 human individuals from eight administrative divisions surrounding the Trijuga forest (Table 1).

### Socio-demographic characteristics of conflict affected people

Males made up the vast majority (91.30%, *n* = 63) of those affected by conflict. Age groups of 40-54 (42.02%, *n* = 29) and 25-39 (36.23%, *n* = 25) were affected more than expected (χ^2^ = 44.406, df = 4, *p* < 0.00001; Table 1). When compared to other ethnic groups, people from Tharu community (36.23%, *n =* 25) and Brahmin & Chhetri (29%, *n* = 20) were involved more in conflicts (χ^2^ = 21.072, df = 4, *p* = 0.0003). The majority (57.97%, *n* = 40) were engaged in farming as their primary occupation and most were illiterate (60.87%, *n* = 42). Those who reported visiting the forest on a daily basis made up the biggest number (71.01%, *n* = 49) of conflict victims than those who visited less frequently (χ^2^ = 79, df = 3, *p* < 0.00001; Table 1).

### Spatial and temporal patterns of conflict

Conflict incidents largely (95.45%, *n* = 63) took place inside the forest apart from a few (*n* = 3) that occurred in plantation areas (*e.g.*, rice field, mango orchard). In 74.24% (*n* = 49) of the incidents visibility condition was poor while remaining had moderate (21.21%, *n* = 14) to good (4.55%, *n* = 3) land cover visibility. Conflicts most typically occurred in narrow gullies (*n* = 21), followed by hill slopes (*n* = 12), riverbank (*n* = 10) and forest trail (*n* = 6), according to location data (*n* = 49) for incidents that occurred in the forest. Conflict incidents mapped using of 4-km^2^ grid cells (*n* = 58) indicated that human-sloth bear conflict is primarily concentrated in the east section of the Trijuga forest (Figure 2).

**Figure 2:**
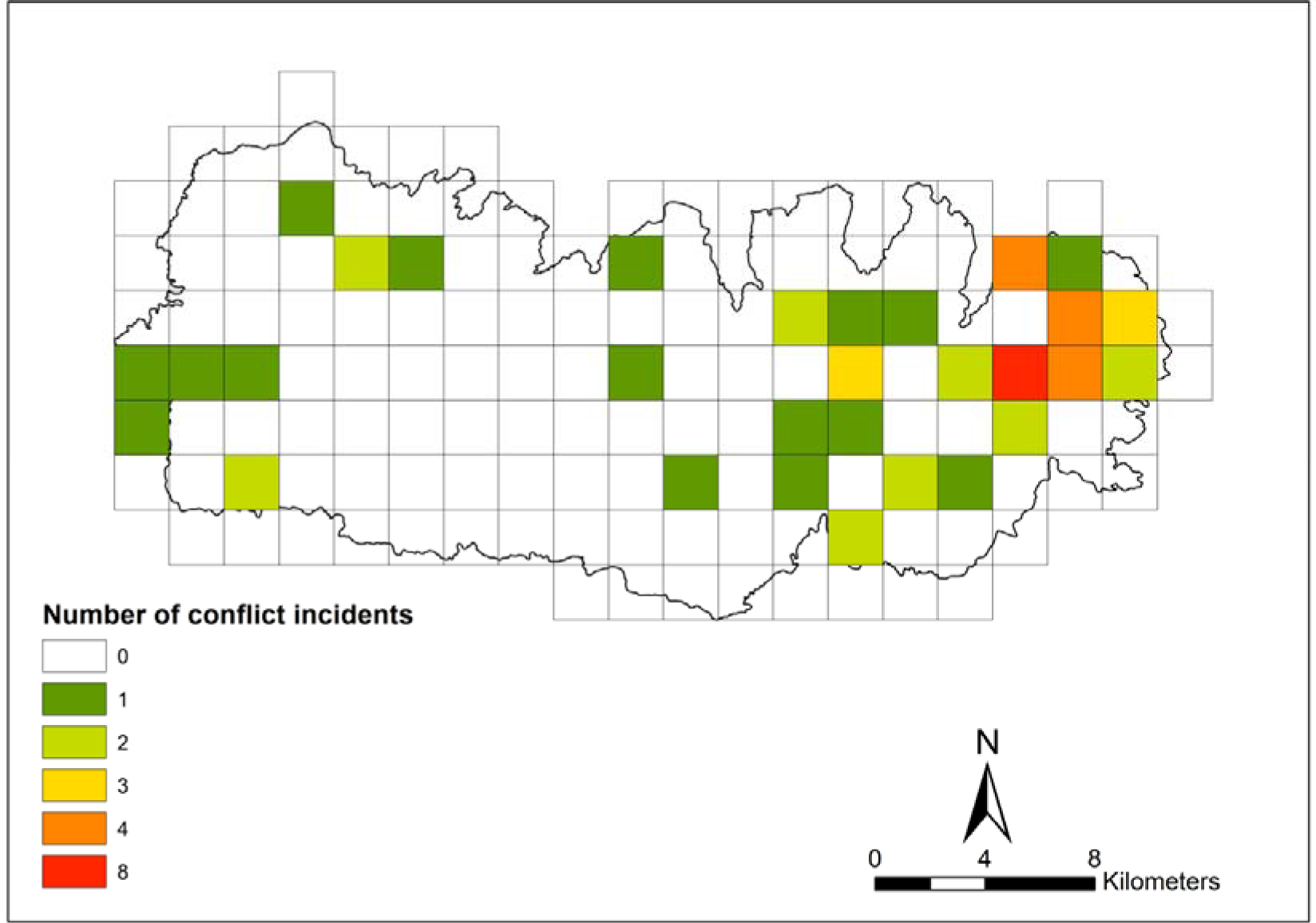
Incidents of human-sloth bear conflict in the Trijuga forest from 1990 to 2021 mapped using 4-km2 grid cells based on the location data (n = 58) provided by the interviewees

On average, 2.06 conflicts were recorded per year from 1990 to 2021 with an attack rate of 1.75. The overall trend of conflict showed an insignificant marginal increment (0.03 *±* 0.02_SE_). When separating the years in three time periods, no significant difference was observed in the frequency of conflicts among them (χ^2^ = 3.90, df = 2, *p* = 0.14), though the trend was found increasing in the most recent decade (Table 3; Figure 3). Conflicts occurred almost throughout the year with no significant variation in terms of months (χ^2^ = 11, df = 10, *p* = 0.35) or seasons (χ^2^ = 3.09, df = 3, *p* = 0.37). Nevertheless, a peak (31.82%, *n* = 21) was observed during the monsoon season, whereas October had the highest number of incidents by month and no incidents were reported on July (Figure 4). Distribution of conflict events was not even throughout the day (χ^2^ = 20.212, df = 4, *p* = 0.0004) and occurred most commonly between 1200 and 1500 (36.36%, *n* = 24; Figure 5).

**Figure 3:**
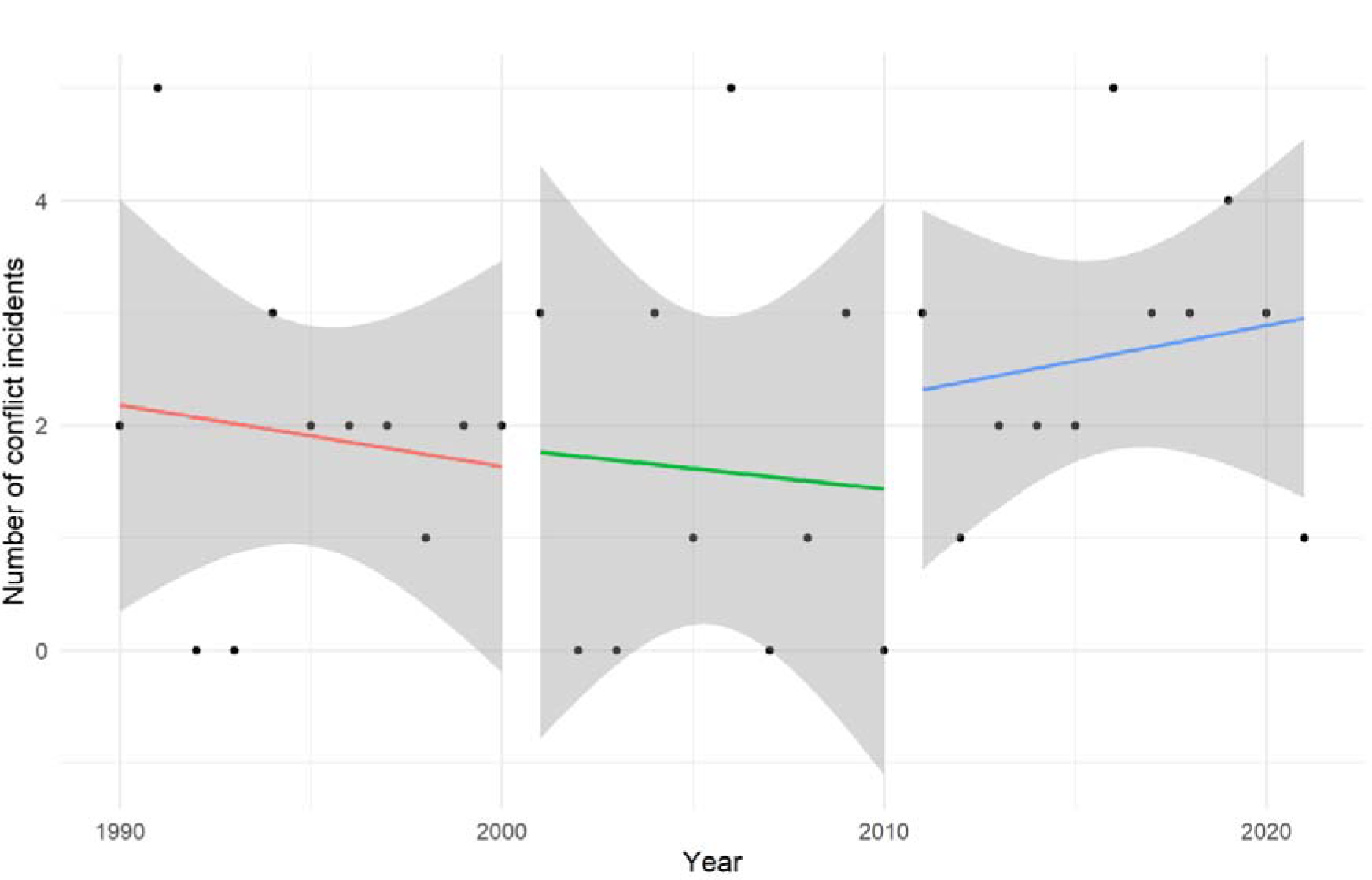
Trend of human-sloth bear conflict in the Trijuga forest across the past three decades

**Figure 4:**
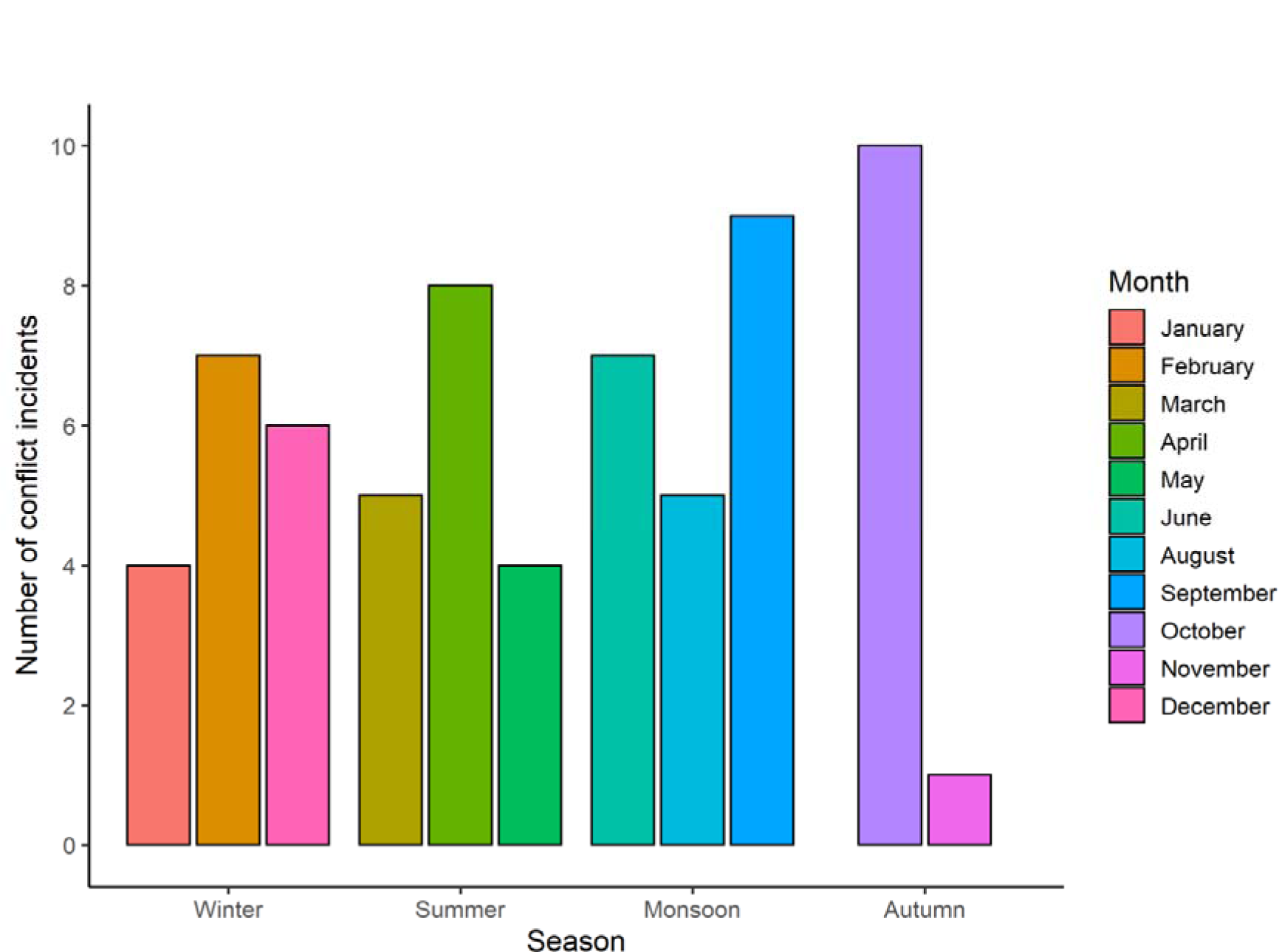
Seasonal and monthly variations in human-sloth bear conflict in the Trijuga forest from 1990 to 2021

**Figure 5:**
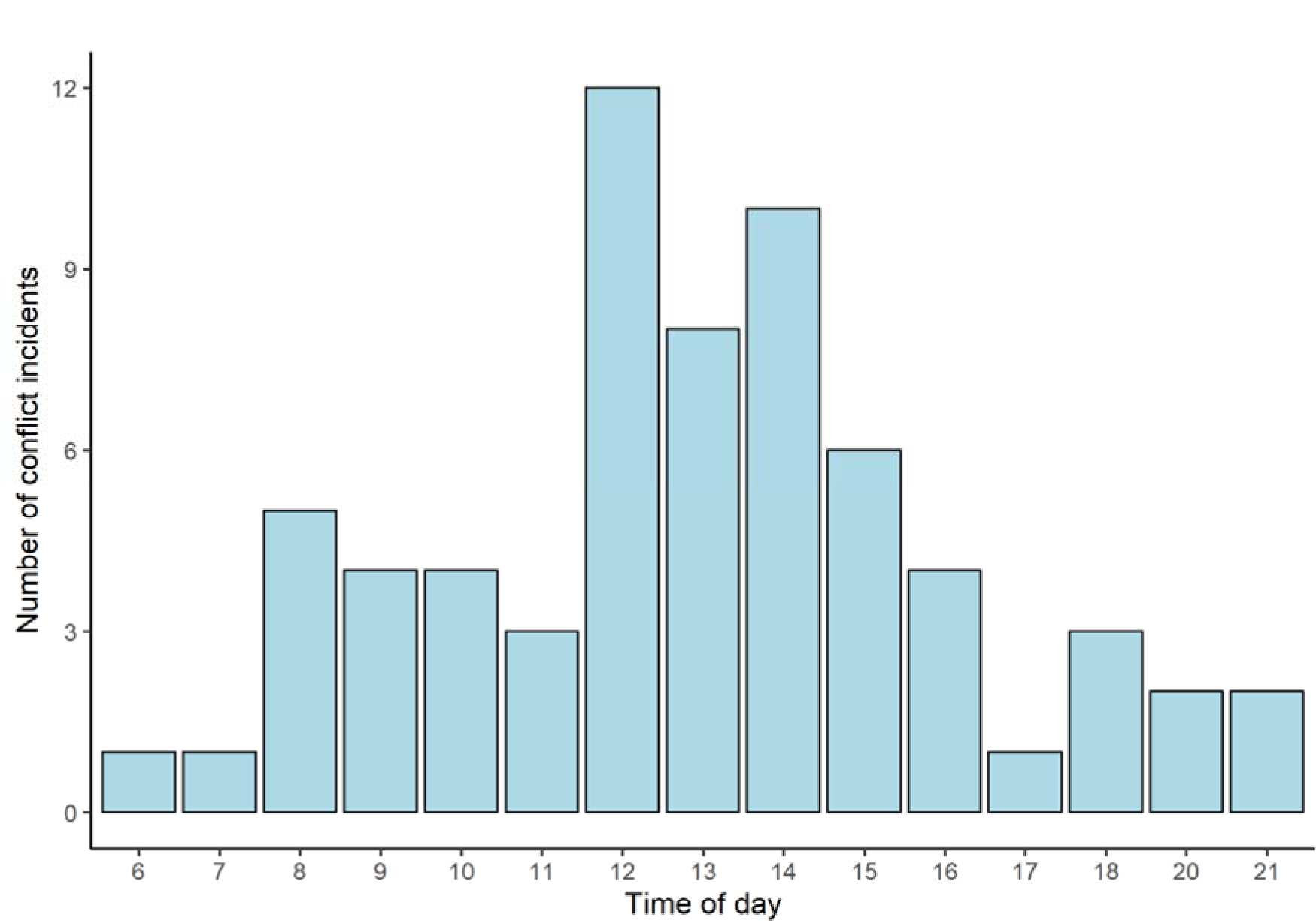
Variation in human-sloth bear conflict by time of the day in the Trijuga forest from 1990 to 2021

**Table 3:** Temporal pattern of human-sloth bear conflict incidents and attacks (mean *±* SD) in the Trijuga forest from 1990 to 2021

| Time period | Average of total conflicts | Average of total attacks |
| --- | --- | --- |
| Annual<sup>a</sup> | 2.06 (1.48) | 1.75 (1.34) |
| 1990-2000<sup>b</sup> | 1.91 (1.38) | 1.91 (1.38) |
| 2001-2010<sup>c</sup> | 1.60 (1.78) | 1.5 (1.58) |
| 2011-2021<sup>b</sup> | 2.64 (1.21) | 1.82 (1.17) |
| Per month<sup>d</sup> | 5.5 (3) | 4.67 (2.14) |
| Per season<sup>e</sup> | 16.50 (4.12) | 14 (4.83) |

<sup>a</sup>total number of years = 32; <sup>b</sup>number of years = 11; <sup>c</sup>number of years = 10; <sup>d</sup>number of months = 12; <sup>e</sup>number of seasons = 4

### Characteristics and circumstances associated with the conflict

Conflict incidents mostly involved multiple bears however; single bears’ involvement was more (46.03%, *n* = 29) when compared separately to female with offspring and similar-sized pairs (Table 3). Apart from the cases where people were unsure about bear activity before encounters, bears were most commonly (26.98%, *n* = 17) reported to be resting, either alone, in pair, or with cubs (Figure 6).

**Figure 6:**
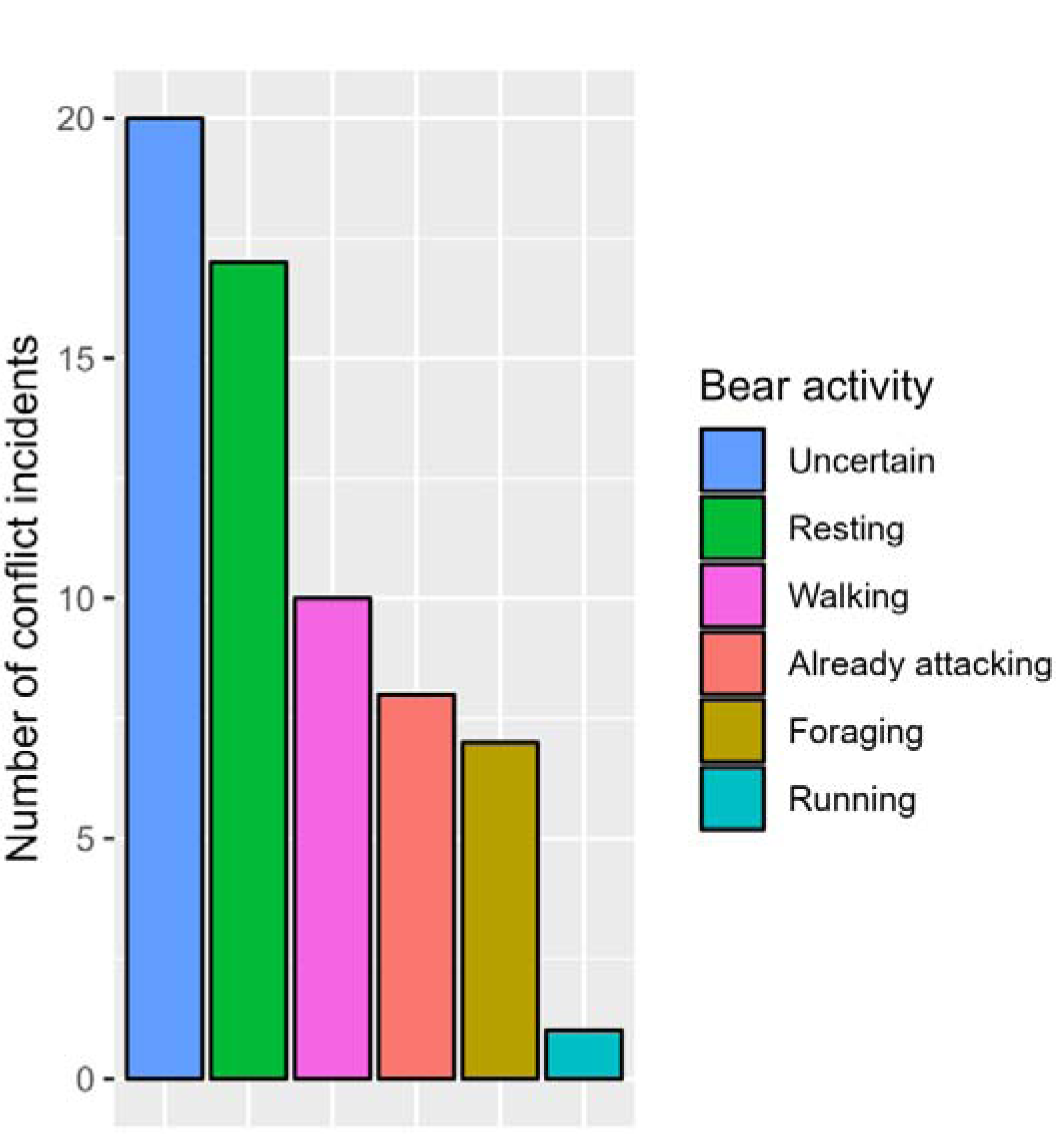
Activity of sloth bears immediately before conflicts in the Trijuga forest

The primary motive for most persons (33.82%, *n* = 23) visiting the site of conflict was to harvest fodder (Figure 7). However, walking was their major activity in a large majority of conflict incidents (64.52%, *n* = 40; Figure 8). They made no talks or noise prior to the encounter in 69.70% (*n* = 46) of cases. There was data on the availability and number of human companions with conflict victims for 64 conflict incidents. There were no companions in half of the events. In 18.75 % (*n* = 12) of the events, there was only one companion, and in 31.25 % (*n* = 20) of the events, there were two or more. In 30.35% (*n* = 17) of the attack episodes, people reported actively fighting with the bear, and they reported hitting the bear once or twice in 16.07% (*n* = 9) of them.

**Figure 7:**
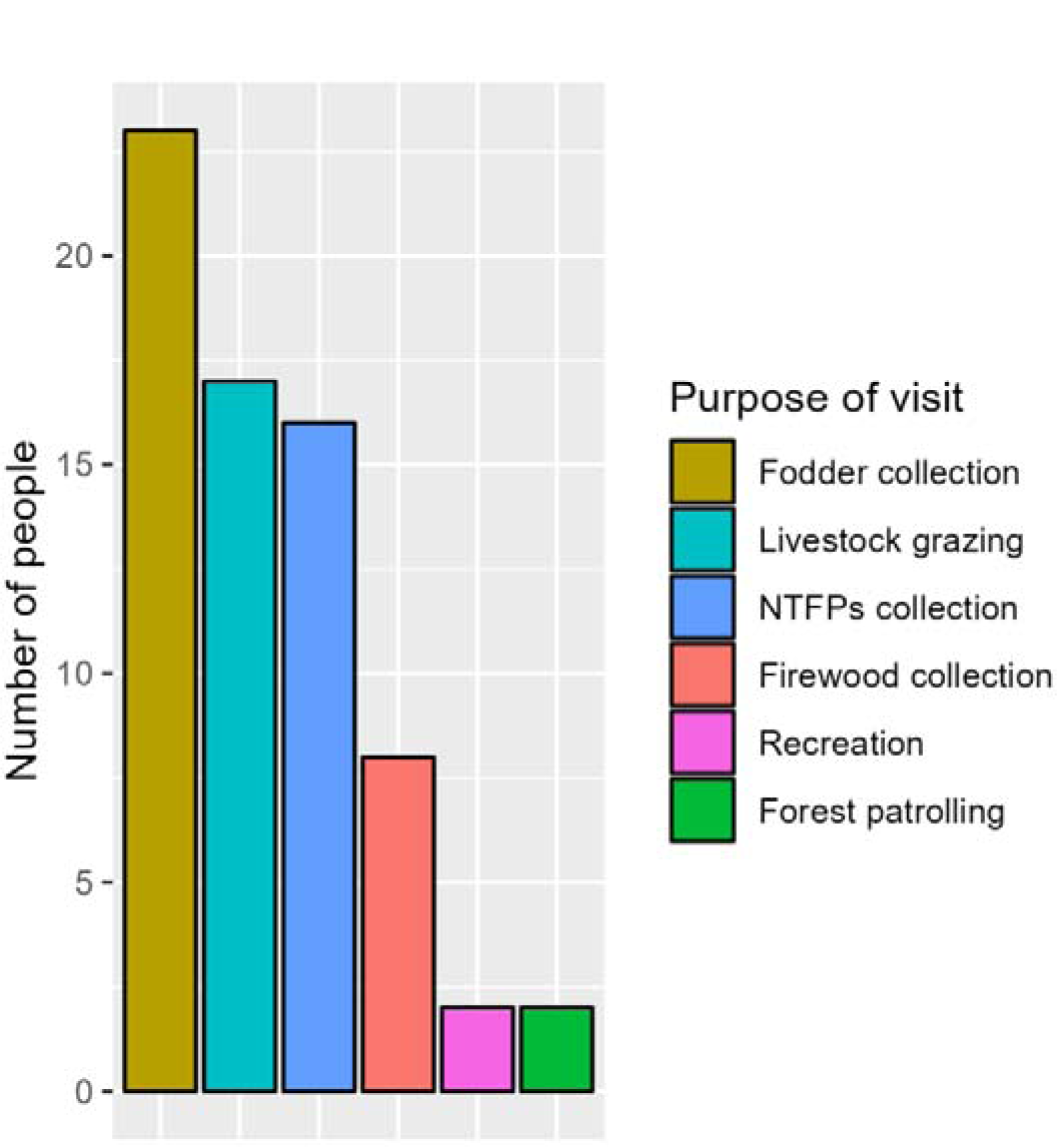
People’s purpose of visiting the site of conflict with sloth bears in the Trijuga forest

**Figure 8:**
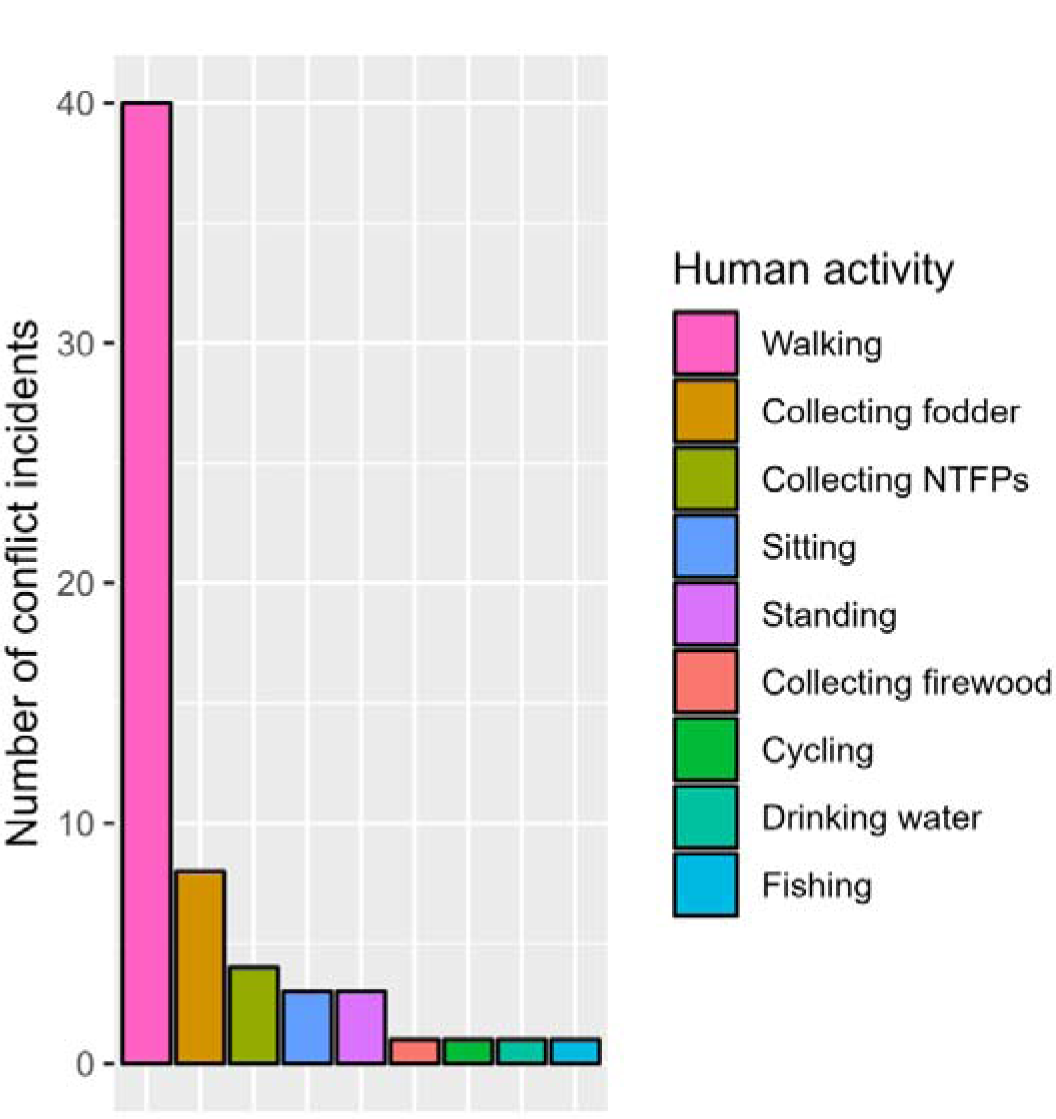
Activity of humans immediately before conflicts with sloth bears in the Trijuga forest

### Injuries resulting from attacks

Humans were injured in 54 out of 56 attack events (Number of humans injured = 59) and bears were injured to some degree in 19 events. The majority (44.07%, *n* = 26) of people received serious injuries from bear attacks (χ^2^ = 13.203, df = 3, *p* = 0.004) compared to moderately serious (25.42%, *n* = 15), moderate (13.56%, *n* = 8), and minor injuries (16.95%, *n* = 10). Of all the people bearing injuries, 8.47% (*n* = 5) died within a few hours of attack, whereas two people died between 1 and 3 years because of complications developed from the wounds. Injuries from bear attacks were most commonly (35.09%, *n* = 40) recorded in the head-neck region of the body. This resulted in people losing either one or both of their eyes causing partial or complete blindness (*n* = 16), loss of scalp (*n* = 14), injuries in ear (*n* = 7), mouth (*n* = 5) and damage of other facial structures (*n* = 17). Other body parts that was injured were hands and arms (28.07%, *n* = 32), legs (19.30%, *n* = 22), chest (6.14%, *n* = 7), back (5.26%, *n* = 6), stomach (5.26%, *n* = 6) and perineum (0.88%, *n* = 1). The observed difference in frequencies among injured body parts was statistically significant (χ^2^ = 60.421, df = 5, *p* < 0.00001). A moderate level of association was found between human group size and level of injuries received (χ^2^ = 10.997, df = 3, *p* = 0.01; Cramer’s V = 0.43), with people having no companions being more likely to receive serious injuries compared to those having one or greater number of companions (Figure 9).

**Figure 9:**
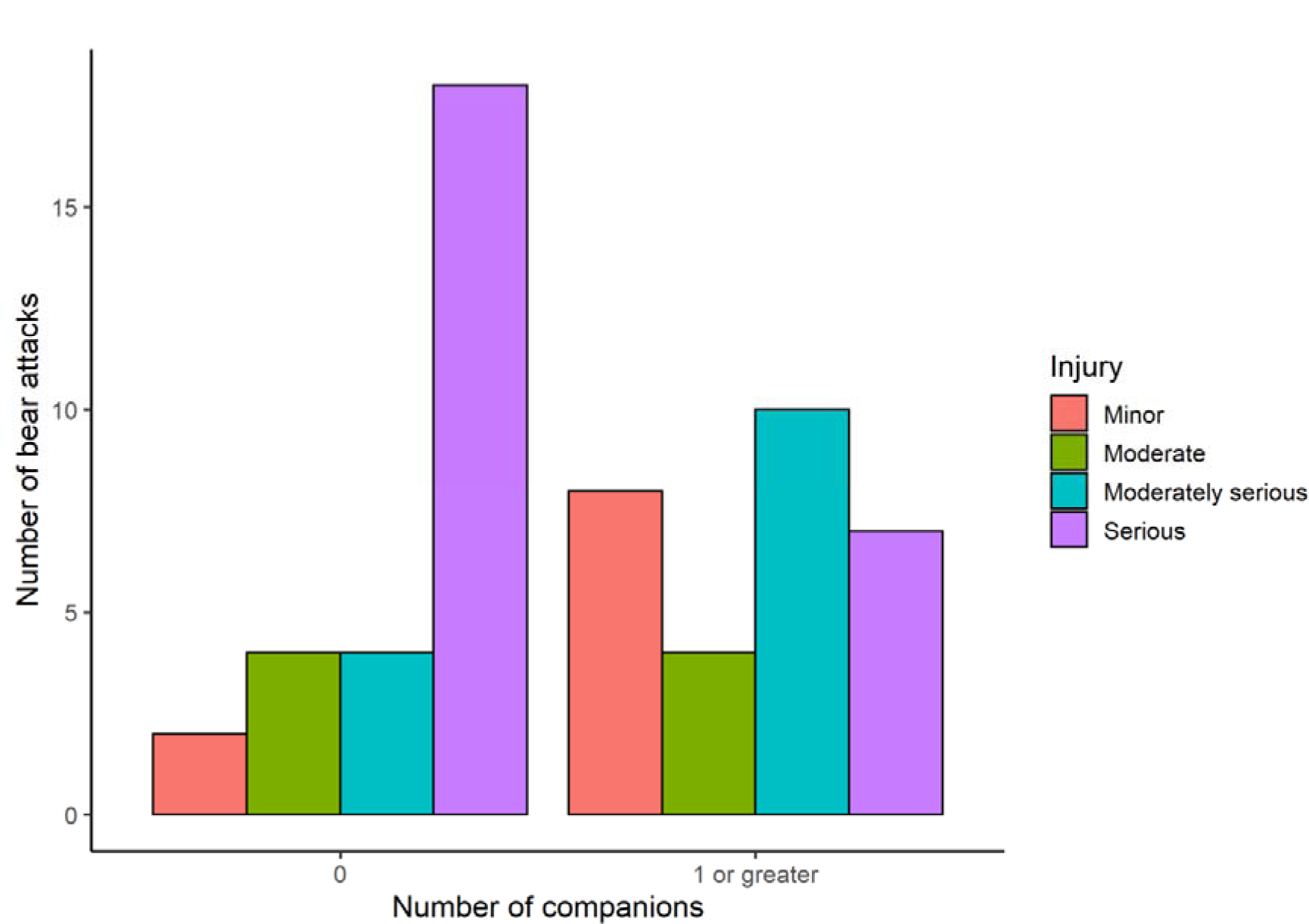
Frequency of sloth bear attacks grouped by number of companions with conflict victims and level of injuries received from attacks

### Predictors of human attacks

The top model for predicting human attacks by bear included an additive combination of human and bear group sizes (AICc weight = 0.194; Appendix 2). However, no single model was dominant and all the four variables were represented in the competing models (models with ΔAICc < 2). Among the four variables, highest cumulative weight was attained by human group size (Summed AICc weight = 0.692), followed by bear group size (0.543), noise (0.243), and poor visibility (0.241). According to model averaged regression coefficients, the size of the bear group and low visibility were shown to be positive predictors of sloth bear attack, while the size of the human group and noise were found to be negative predictors (Table 5). The influence of visibility was strong and significant, with the 95% CIs not containing 0, while the CIs for human group size marginally overlapped 0. CIs of the remaining two variables more strongly overlapped 0 (Table 5).

**Table 4:** Social structure of bears engaged in conflicts with humans in the Trijuga forest

| Social structure of bears | Number of incidents ($n = 63$) |
| --- | --- |
| Single bear | 29 |
| Female with cubs | 23 |
| Similar sized pair | 11 |

**Table 5.** Weighted β estimates, standard errors, and 95% confidence intervals of the variables devised to test influence on sloth bear attack probability

| **Variables** | **$\beta$** | **$SE(\beta)$** | **LCI** | **UCI** |
| --- | --- | --- | --- | --- |
| (Intercept) | 1.7861 | 0.6214 | 0.5682 | 3.0040 |
| Bear group size | 0.5467 | 0.4372 | -0.3102 | 1.4036 |
| Human group size ($\leq 50m$) | -0.2542 | 0.1554 | -0.5588 | 0.0504 |
| Making noise (Yes) | -0.4693 | 0.767 | -1.9726 | 1.0340 |
| Poor visibility (Yes) | 2.5803 | 0.8066 | 0.9994 | 4.1612 |

## Discussions

Our findings are comparable with previous studies from developing countries in demonstrating the heightened susceptibility of forest-dependent rural communities to wildlife conflicts, including with sloth bears [17,20,34,44,45]. There were considerable differences across different socio-demographic categories (e.g., ethnicity, age, education, gender, and occupation) in their frequency of being involved in conflicts. These suggested that the most vulnerable groups are working-age men who practice forest-based livelihoods (e.g., farming), have little to no formal education, and frequent the forest almost daily for activities like resource gathering and livestock grazing. Throughout southern Nepal, people who have experienced human attack-associated conflicts with other large mammals such as tigers, elephants, and leopards are also reported to share similar socio-demographic characteristics [26,29,30]. This indicates that conflict management approaches tailored to meet the socioeconomic concerns of the vulnerable groups identified by this study can promote coexistence with all the major species in this part of Nepal.

We reported a considerable spatial difference in conflict incidences within our study area. Conflicts were mostly concentrated in the east section of the Trijuga forest and were relatively fewer in other parts (Figure 3). Our recent study documented higher space use probability of sloth bears in this part of the forest which was attributed to the availability of important ecological resources for bears (e.g., termite mounds and water sources) [36]. Because human activities were also significant in this part [36], the greater overlap in sloth bear and human use of the forest could have contributed to a high number of human-bear conflicts. There are reports of higher chances of human-wildlife confrontations in areas where spatial overlap or proximity between humans and wildlife is more [46,47].

In many human-dominated and relatively disturbed natural areas, conflicts with sloth bear have frequently been reported within or close to human usage areas such as agricultural fields and settlements [17,21,23]. However, around Trijuga forest conflicts predominantly happened inside forests despite it being surrounded on most sides by human modified areas. Forest cover within the present study area has remained relatively stable over the years, despite a considerable decline in forest cover across the Terai and Churia region, especially outside protected areas [48]. As studies have shown that sloth bear occurrence is positively associated with forest cover [49,50], we speculate that the availability of comparatively stable forest cover in this area may have benefited in meeting sloth bear habitat needs to a certain extent [51], reducing their presence in areas of human use.

Our study inferred that poor land cover visibility significantly elevates the possibility of a sloth bear attack if encountered. Previous researches on human-bear conflict have come up with similar conclusions [9,24], albeit no empirical testing similar to ours was undertaken to substantiate the findings. In our case, most human-bear confrontations took place in areas with rough terrain and dense vegetation (e.g., narrow gullies, hill slopes, and thickets of riverbanks). Sloth bears are known to prefer similar areas for resting, especially during the day, and such places also provide ideal feeding habitats to sloth bears [51]. In the Trijuga forest similar observations have been made based on the indirect signs of sloth bears [36]. Given these insights into sloth bear ecology and our study’s findings that resting and foraging were important known activities of sloth bears at the time of encounter, it is reasonable to conclude that people mostly surprised bears at a close range in areas of poor visibility when bears were at resting or feeding sites [20]. In this context, habitat features associated with poor land cover visibility (e.g., high vegetation productivity and terrain ruggedness) might be important determinants of sloth bear attacks in this area, as documented elsewhere [52].

In investigating the temporal patterns of sloth bear conflicts, we did not observe statistically discernible change over the past three decades. However, the rate of conflicts in the most recent decade was found increasing at a rate greater than the annual average from 1990 to 2021. The seasonal or monthly differences in conflict incidents involving sloth bears were also not supported statistically by our study. It might be because human activities in this area are somewhat consistent throughout the year. Nevertheless, our documentation of more conflicts (48%) in the monsoon to post-monsoon season is comparable with findings from eastern and central India [21,23]. During the monsoon and post-monsoon, the vegetation productivity in the forests of Churia region is greatly increased. A higher number of conflicts may have been observed during this time of the year due to the decreased visibility caused by the increased foliage cover [21]. There may also be an increase in individuals visiting the forest to collect non-timber forest products (NTFPs), such as bamboo sprouts, mushrooms, and fiddleheads at this time of the year.

Interestingly, however, we were unable to document any conflicts in July. In southern Nepal, paddy cultivation is at peak during July [53], and since paddy is the most important crop for the locals, they are most likely to be working in the fields during this time of year. As an outcome, there may have been lower regularity of people visiting the forest leading to minimal human-sloth bear encounters. Throughout the sloth bears’ range, monthly variations in the frequency of conflicts dependent on the level of human activity within or close to bear habitat have been noted [17,20,24]. The same explanation should also be applicable in describing the influence of time of the day on conflict with sloth bears. Conflicts predominantly took place between 0900 and 1500 (65%) coinciding with the time when humans tend to be more active in the forest in our study area.

Our documentation of slightly greater involvement of multiple bears in conflicts varies from the majority of earlier studies on sloth bears that report single bears to be generally engaged in such situation [17,18,20,23]. The regression analysis further suggested that, in our situation, probability of an attack rises as bear group size increases. The observed inference may be explained by analyzing the sociobiology of sloth bears. For instance, even though sloth bears are solitary in nature, family grouping of females with cubs is common, and to a lesser extent, coalitions of sub-adults have also been observed [12,15]. It has also been noted that during mating season, multiple male sloth bears may congregate around receptive females [12]. When with cubs, females are highly defensive, making them more prone to attack intruders [15]. Similarly, aggressive behavior and fights between males have indeed been documented during mating sessions, whereas the coalitions of young bears is suspected to be a tactic for defense against potential predators or adult male conspecifics [12,14]. Therefore, encounters with humans in either scenario may also increase the bears’ propensity to attack. Moreover, since our study could not derive a strong support for this evidence, as indicated by the 95% CIs overlapping 0, future studies are essential to support this hypothesis and gain more robust insights in this matter.

Studies other than ours have also provided evidence in favor of the effect of human group size in predicting the outcome of human-large carnivore encounters, including with bears [9,10]. In general, it is implied that humans in groups are less vulnerable to attacks by large carnivores like bears and that individuals acting alone are more likely to be involved in attacks that result in fatalities or serious injuries [9,10,51]. Not only did we find a higher number of solitary persons encountering sloth bears, but we also observed a link between human group size and the severity of injuries, with single people suffering more serious injuries, and all fatal attacks involved solitary persons. Serious injuries by sloth bear attacks were almost always due to the wounds sustained in the head and neck region of the body. Despite reports of sloth bears inflicting similar patterns of injuries in other range areas [18,20,23], the proportion of victims in our case who suffered severe injuries (44.07%) was higher compared to other studies [17–19]. The documented fatality rate (8.47%) was also among the highest for sloth bears reported to date [19,21,23,51]. This severity observed in our study may be the result of human behavior during conflicts. For example, in around 46% of bear attack cases respondents in our study mentioned fighting or at least frightening the bears. It is noted that such conduct can make bears feel more threatened putting them under more pressure to engage in attack for longer periods of time when their natural instinct is to leave the area unharmed [17–19].

## Conclusions

A larger proportion of sloth bear habitat in Nepal is unprotected, where they are forced to share space with humans, often giving rise to human-sloth bear conflicts. Recognizing that these shared habitats are important for both humans and sloth bears, conflict mitigation should address the issues of both parties. Our study inferred that human-sloth bear conflict in the studied region follows a predictive pattern associated mainly with habitat and human-related factors. Higher rates of overall conflicts as well as attacks were in areas having poor visibility conditions and in cases where humans were alone. Hence, it is imperative to understand that effective human-sloth bear conflict management would first require the identification of conflict-risk habitats and then carefully regulating human activities and behaviors in those areas. Outside Nepal’s protected areas, where forests are mostly managed by communities, local forest authorities and community members can work together to map such risky habitats and, if possible, monitor sloth bear movements through periodic surveys. This will not only help minimize direct conflicts but could also improve local people’s ability and attitude to coexist with sloth bears. Further, our study emphasizes the importance of community education programs to better inform the locals about sloth bear ecology and behavior, precautions to respond to bear attacks, and protecting vital body parts during attacks.

## Acknowledgements

Financial support for this study was provided by the International Association for Bear Research and Management (IBA-SG 2020/21-03: fRI Fellowship). We would like to acknowledge IDEA WILD for providing equipment support. Research permit was provided by the Department of Forests and Soil Conservation of the Government of Nepal (Ref no. 134/078/079). We are sincerely thankful to the people of forest offices (Division, sub-division and community forests), local administrative offices, and local communities surrounding the Trijuga forest for their valuable help in identifying conflict affected people. We are indebted to the local field assistant Mr. Amit Kumar Chaudhary and conflict victims from around the Trijuga forest for their support during data collection. We are grateful to Prof. Karan Bahadur Shah, Dr. Prakash Kumar Paudel, and Mr. Yadav Ghimirey for their support in securing project funding.

