## Appendix 1 for "Patterns and predictors of human-sloth bear conflict in Nepal: Implications for coexistence in unprotected forest landscapes"

**Human-sloth bear conflict assessment in the Trijuga forest**

### ***Questions for assessing conflict***

**1. Temporal data of conflict:**

a. Year_______________ b. Month_____________ c. Time______________

**2. Approximate geographic location (Show map)**

___________________________________________________________________________

**3. Land cover visibility (Mention the vegetation and terrain features):**

a. Poor_________________ b. Moderate________________ c. Good__________________

**4. Human activity during encounter:**

___________________________________________________________________________

**5. Bear activity during encounter:**

___________________________________________________________________________

**6. Encounter result into attack?** a. Yes b. No

**7. Human response to encounter or attack:**

a. Ran away b. Climbed a tree c. Hid behind something d. Threw stones or weapons at bear d. Shouted e. Fought with bear f. Laid on the ground c. Slowly stepped back

Details_____________________________________________________________________

__________________________________________________________________________

**8. Bear response during encounter or attack:**

a. Vocalized b. Ran away c. Stood up on hind legs d. Approached slowly e. Rapidly charged f. Chased for some time and left g. Used claws h. Used teeth

Details_____________________________________________________________________

___________________________________________________________________________

**9. Number of human companions nearby (Mention approximate distance):**

___________________________________________________________________________

**10. Response of companions during encounter or attack:**

________________________________________________________________________

**11. Demographics of bears involved in attack or encounter:**

________________________________________________________________________

**12. Consequences of attack or encounter:**

a. Inflicted injuries to human b. Bear was injured c. Bear was killed d. Bear ran away

**Details**__________________________________________________________________

________________________________________________________________________

**13. Type of injuries inflicted to humans:**

________________________________________________________________________

________________________________________________________________________

________________________________________________________________________

### ***Questions for assessing socio-demographic information***

**14. Full name: _______________________ 15. Age: _____ 16. Gender: _________**

**17. Ethnicity_____________18. GPS location: ___________________________________**

**19. Family size: ____________ 20. Major occupation: ________________**

**21. Municipality/village: ____________________________________________**

**22. Frequency of forest visit: _________________________________________**

**23. Education: ___________________________________________**

**24. Lived in this area?**

a. Since childhood b. >20 years c. 10-20 years d. 5-10 years e. <5 years

**25. Major purpose of forest visit**

________________________________________________________________________

________________________________________________________________________
