## Appendix 2 for "Patterns and predictors of human-sloth bear conflict in Nepal: Implications for coexistence in unprotected forest landscapes"

Table 1: List of models built using the firth's bias-reduced logistic regression for determining the influence of variables on the probability of a sloth bear attack in the Trijuga forest

| Models | df | logLik | AICc | ΔAICc | AICc weight |
| --- | --- | --- | --- | --- | --- |
| Bear group size + human group size | 3 | -23.552 | 53.5 | 0 | 0.194 |
| Human group size | 2 | -24.893 | 54 | 0.47 | 0.154 |
| Human group size + making noise (yes) | 3 | -24.346 | 55.1 | 1.59 | 0.088 |
| Bear group size | 2 | -25.509 | 55.2 | 1.7 | 0.083 |
| Bear group size + human group size + poor visibility (yes) | 4 | -23.285 | 55.3 | 1.75 | 0.081 |
| Bear group size + human group size + making noise (yes) | 4 | -23.343 | 55.4 | 1.87 | 0.076 |
| Human group size + poor visibility (yes) | 3 | -24.616 | 55.6 | 2.11 | 0.068 |
| Null | 1 | -26.996 | 56.1 | 2.54 | 0.055 |
| Making noise (yes) | 2 | -26.066 | 56.3 | 2.82 | 0.048 |
| Bear group size + making noise (yes) | 3 | -25.073 | 56.6 | 3.04 | 0.042 |
| Bear group size + poor visibility (yes) | 3 | -25.249 | 56.9 | 3.39 | 0.036 |
| Bear group size + human group size + poor visibility (yes) + making noise (yes) | 5 | -23.075 | 57.2 | 3.7 | 0.031 |
| Poor visibility (yes) | 2 | -26.733 | 57.7 | 4.14 | 0.025 |
